# *HoloBio*: A Holographic Microscopy Tool for Quantitative Biological Analysis

**DOI:** 10.64898/2026.01.20.700497

**Authors:** Waira Mona, Maria J. Gil-Herrera, Emanuel Mazo, Daniel Córdoba, Sofia Obando, Maria J. Lopera, Rene Restrepo, Carlos Trujillo, Ana Doblas, Raul Castaneda

**Affiliations:** SOPHIA Research Group Optics and Photonics Laboratory, School of Applied Science and Engineering, Universidad EAFIT, Medellín, Colombia 050022; ECE Department, University of Massachusetts Dartmouth, New Bedford, Massachusetts, 02747, USA

## Abstract

Holographic imaging in microscopy enables label-free quantitative information of biological specimens and has found applications across a wide range of biomedical studies, from cell morphology to particle dynamics; yet its widespread adoption is often limited by the lack of accessible and standardized analysis software. We present *HoloBio*, an open-source, Python-based graphical user interface developed to address this issue. This software offers two primary operational modes: a *Real-Time* mode that enables live processing of holograms at video frame rates, and an *Offline* mode designed for post-processing previously recorded holograms. *HoloBio* is compatible with holograms recorded using both lens-based and lensless systems, supporting off-axis architectures in telecentric and non-telecentric configurations, as well as slightly off-axis and in-line optical setups. The software incorporates tools for cell tracking, phase profiling, thickness estimation, and morphological analysis, including cell counting and object area quantification. *HoloBio* is designed to be accessible for users without coding expertise, offering a reproducible, high-throughput environment tailored for researchers in biology, biophotonics, and biomedical imaging.

## Introduction

Holographic imaging in microscopy has become a key label-free, video-rate quantitative imaging technique for studying biological specimens because holograms encode both the amplitude (absorption/attenuation contrast) and the quantitative phase (optical path length linked to refractive index and thickness) of transparent samples(1). This capability enables rich, non-invasive characterization of biological samples such as long-term live-cell monitoring and growth assessment (2), quantitative analysis of cell dry mass and thickness changes(3), and automated characterization of morphology and dynamics, including motility and migration trajectories(4). These imaging techniques can also support volumetric inspection and 3D tracking of motile cells and microorganisms, both in lens-based Digital Holographic Microscopy (DHM) and in lensless or in-line configurations (DLHM/DIHM)(5). The comprehensive quantitative information encoded in DHM and DLHM, spanning optical path length, refractive index, thickness, and morphology, has culminated in a growing body of translational applications, enabling objective, label-free assessment of pathological changes in unstained specimens (6).

This versatility has driven the development of a broad ecosystem of reconstruction and analysis tools across multiple programming environments (7–13) [Refs]. In the open-source domain, several libraries provide end-to-end holographic processing capabilities, including HoloPy, which supports hologram reconstruction and light-scattering workflows in Python(9), and pyDHM, which implements phase-shifting and phase-compensation pipelines for multiple DHM configurations (8). At the numerical propagation level, reusable computational backends form the foundation of many DHM and DLHM reconstruction pipelines. For example, CWO++ provides CPU and GPU-accelerated diffraction and propagation routines (12), whereas JDiffraction offers Fresnel and angular-spectrum propagation methods (10). Beyond core propagation and reconstruction, additional projects address higher-level holography workflows. OpenHolo supports hologram generation, reconstruction, and signal processing (13). DHM capabilities have also been integrated into widely adopted biomedical imaging platforms through ImageJ-based toolsets, including HoloJ and other DHM-focused plugins for simulation, reconstruction, and analysis workflows (11,14). Similar ImageJ-based approaches have been extended to lensless and in-line holographic configurations(15). Complementing these open-source efforts, commercial platforms, including Holo4D and other proprietary DHM software suites (16,17), provide fully integrated reconstruction and analysis environments, underscoring the maturity and broad adoption of digital holographic microscopy across both research and translational settings.

However, despite their technical contributions, existing tools present important limitations that hinder broader adoption. Many libraries require advanced programming expertise and a strong background in Optics, thereby limiting their accessibility to researchers in biological and clinical environments. Plugin-based solutions, while widely popular in the biomedical imaging community, tend to be task-specific and often lack cohesive integration across hologram acquisition, reconstruction, and downstream biological quantification. Commercial software platforms offer robust and polished functionalities, but their closed and proprietary nature limits accessibility and widespread use in academic research. As a result, a significant usability gap still remains for biomedical researchers who could otherwise benefit from the versatility of DHM and DLHM but are impeded by the complexity and fragmentation of existing software ecosystems. The lack of a unified, biology-oriented graphical user interface continues to constrain the broader adoption of DHM and DLHM in translational applications such as cell biology and histopathology, where extracting quantitative phase and morphological information from label-free samples holds significant scientific and clinical relevance.

To address these challenges, we present *HoloBio*, a free and open-source Python-based graphical interface software specifically designed for biological applications using DHM and DLHM. *HoloBio* supports a broad range of experimental configurations and integrates multiple reconstruction strategies tailored to different optical geometries, including in-line, slightly off-axis, and off-axis holography. The software is compatible with for both real-time (video-rate) data acquisition and the processing of previously recorded holograms. Beyond hologram reconstruction, *HoloBio* provides a suite of biology-oriented tools aligned with standard microscopy workflows, including micrometric image scaling, digital refocusing via numerical propagation, quantitative phase mapping, thickness estimation, phase profiling and particle tracking. In addition, the platform also features biological analysis utilities such as particle and cell counting, morphology assessment, and speckle noise reduction, enabling robust and reproducible interpretation of label-free imaging data.

## Design and Implementation

*HoloBio* is structured around two main operational modes: (1) *Real-Time* Hologram Processing and (2) *Offline* Hologram Processing. Each mode is subdivided into two specialized packages corresponding to the holography modalities: Digital Holographic Microscopy (DHM) (1) and Digital Lensless Holographic Microscopy (DLHM) (18). This modular organization ensures that each functional interface is optimized for a specific optical configuration and processing workflow.

### Initialization and Mode Selection

Upon launch, HoloBio presents a graphical welcome interface that guides users to intuitively select the operational mode and digital holography modality (**Fig. *1***). Depending on the selected modality, the software dynamically loads the corresponding interface, optimized for either live acquisition or reconstruction of previously recorded holograms.

**Fig. 1.**
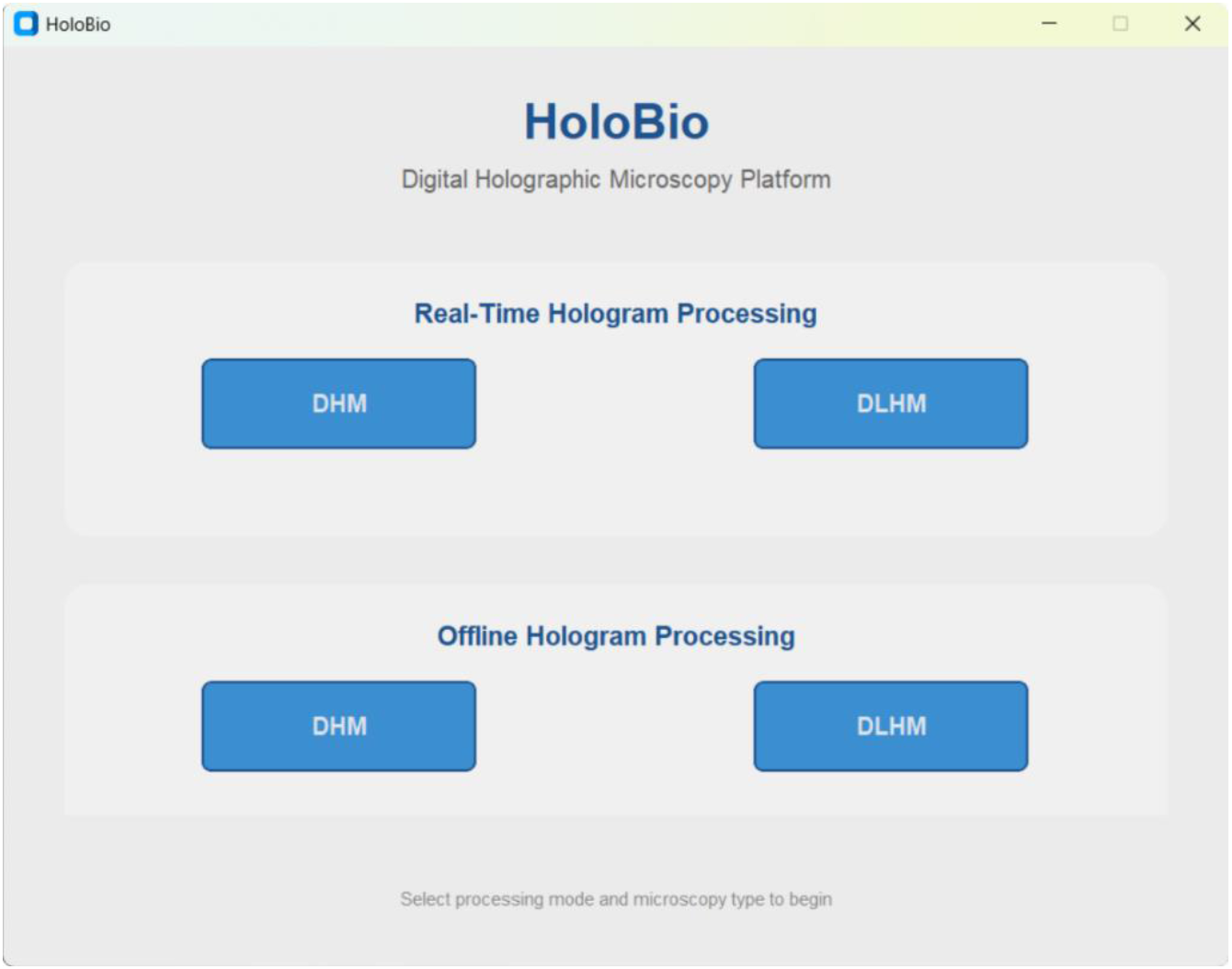
Startup interface of *HoloBio*. The welcome screen allows users to select between *Real-Time* or *Offline* processing modes and choose the desired microscopy modality (DHM or DLHM).

### *Real-Time* Hologram Processing

This mode allows users to live-acquire and reconstruct holograms using a digital camera connected to the system. For both DHM and DLHM imaging modalities, the software provides real-time video-rate visualizations of amplitude and phase-compensated reconstructed images. In this package, users can also import previously recorded hologram videos for reconstruction, as well as amplitude or phase images for particle tracking. Additionally, individual frames can be processed for quantitative phase analysis, speckle noise reduction and conventional image enhancement.

#### *Real-Time* DHM package

This package is designed for DHM systems operating in off-axis configuration under telecentric regime (19). It provides a live interface for real-time visualization of raw holograms alongside their corresponding reconstructed amplitude and phase images. These reconstructed images are computed frame-by-frame using the semi-heuristic phase compensation (SHPC) method (20). Additional features include real-time display of the hologram Fourier Spectrum with interactive zoom, allowing inspection of specific regions in spatial or frequency domain. This capability is particularly useful for system alignment and fine-tuning of telecentric conditions. Both raw holograms and reconstructed amplitude or phase images can be recorded as video sequences. Additionally, previously recorded hologram videos can be imported for real-time reconstruction, supporting both live and retrospective analysis.

The *Real-Time* DHM package interface is structured into three sections (**Fig. *2***). The top control panel (1) contains primary navigation and management tools. The parameter panel (2) allows users to define the physical parameters required for real-time reconstruction, including the wavelength and the pixel pitch along the X and Y directions. In addition, it provides a Fourier transform (FT) visualization module in which the spatial filter geometry (e.g., circular or rectangular) can be selected, and users may display either the filtered or unfiltered Fourier spectrum in real time. *Compensation Controls* are provided to execute real-time reconstruction, run continuous acquisition, or stop the process. The *Record Options* allow recording sequences of holograms and reconstructed amplitude or phase images. Additionally, the *Particle Tracking* options allow users to upload previously recorded holograms and amplitude or phase videos, generate two-dimensional particle tracking, obtain frame-by-frame particle positions, and save the resulting information. The visualization panel (3) displays the hologram or its Fourier Spectrum on the left and either the phase or amplitude reconstruction on the right, both with zoom and view adjustment capabilities. For a detailed description of the *Real-Time* DHM package and its functionalities, readers are referred to the ***HoloBio* User Manual**.

**Fig. 2.**
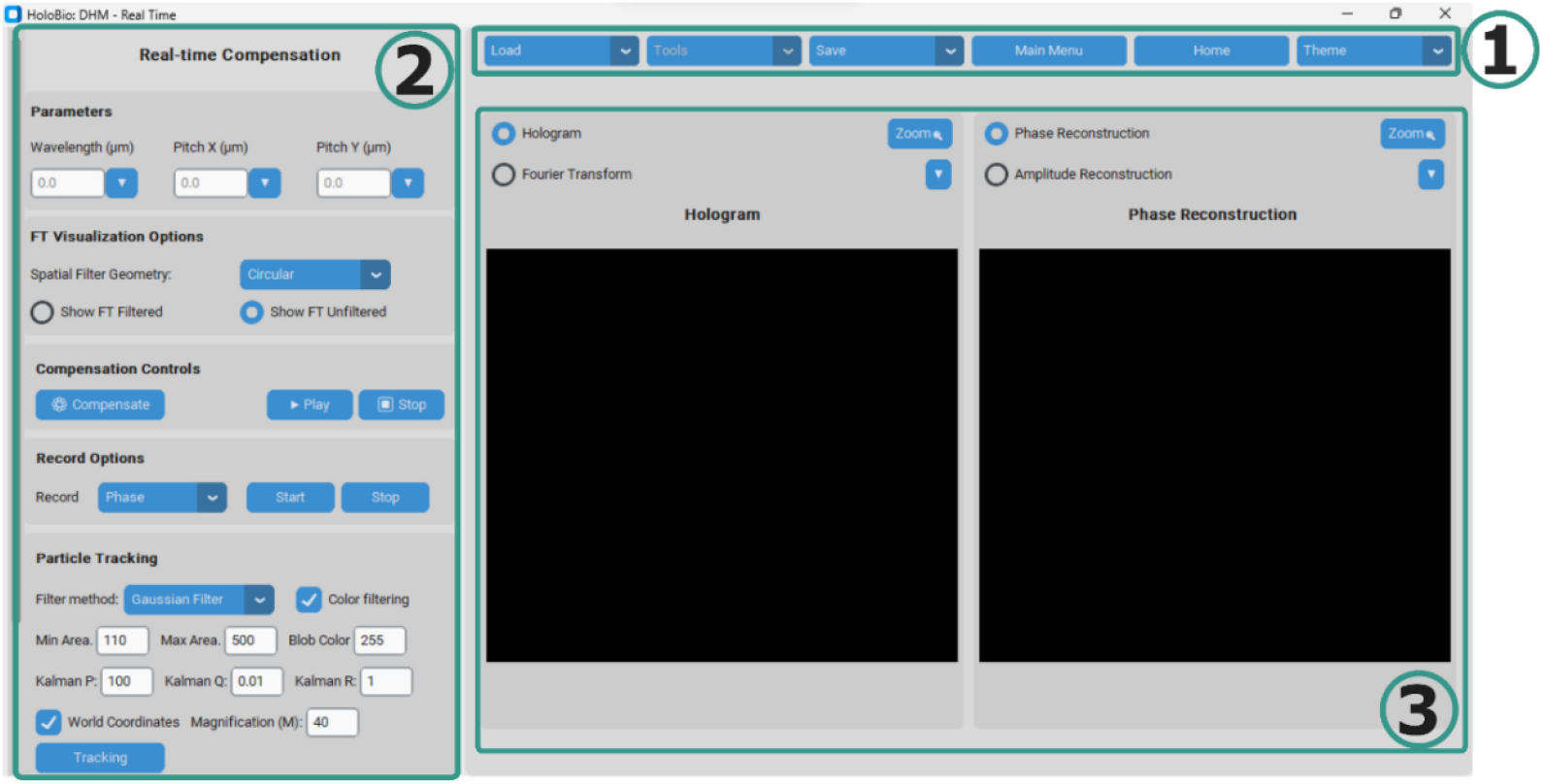
Interface for the *Real-Time* DHM package in *HoloBio*. The interface is organized into three components: (1) the control panel with core functionalities, (2) the real-time processing panel, which provides options for parameter configuration (wavelength, pixel pitch), Fourier transform visualization settings, compensation controls, recording options, and particle tracking tools. (3) The visualization panel for hologram and reconstruction display.

#### *Real*-*Time DLMH* package

This package is intended for DLHM systems employing either spherical (Point-source DLHM) (18) or planar (On-Chip Lensless systems) (21)illumination. It supports real-time visualization of holograms and live reconstruction, providing amplitude and phase outputs. Users can select among three computationally-efficient reconstruction algorithms: the Angular Spectrum approach for plane-wave propagation(22), a discrete version of the Kirchhoff–Helmholtz diffraction integral accounting for spherical wavefronts (KHDI) (15), and a modified Angular Spectrum approach for spherical illumination (MAS) (23). A distinctive feature of this package is the interactive control of the propagation distance, enabling real-time adjustment to achieve optimal focus.

### *Offline* Hologram Processing

The *Offline* mode is dedicated to the numerical reconstruction and analysis of previously recorded holograms. Upon selection, users choose between DHM and DLHM modalities, after which the software launches a specialized interface tailored to the corresponding optical configuration.

#### *Offline* DHM package

This package offers three reconstruction workflows: *Phase-Shifting, Phase Compensation*, and *Numerical Propagation*. The *Phase-shifting* workflow supports six algorithms to reconstruct in-line or slightly off-axis holograms, including the 5-Frames (24), 4-Frames(25), 3-Frames (24), and Quadrature (26) methods, as well as two blind approaches for unknown phase shifts: Blind 3 Raw Frames (27) and Blind 2 Raw Frames (28). The *Phase Compensation* workflow provides four methods for off-axis holograms: Semi-Heuristic Phase Compensation (SHPC) (20), Tu-DHM (29), Non-Telecentric reconstruction (30), and Vortex-Legendre fitting (31). Finally, the *Numerical Propagation* workflow enables reconstruction at arbitrary propagation distances for experimental or simulated wavefields using either the Angular spectrum method or the paraxial Fresnel Transform (22).

Selecting this package opens the corresponding interface shown in **Fig. *3***, which is designed to guide the user through the hologram reconstruction workflows described previously. The interface comprises three main components. At the top of the interface, the **main control panel (1)** provides access to core software functionalities. This includes buttons for loading and saving data, accessing additional tool panels (*Tools*), navigating to the main menu or home screen, and switching between interface themes. The **left panel (2)** provides access to the available DHM processing methods: *Phase Shifting, Phase Compensation*, and *Numerical Propagation*. Depending on the selection, the control panel dynamically adapts to display relevant parameters and tools for the chosen method. The **visualization panel (3)** serves as the main visualization area, where users can display the original hologram, its Fourier transform, and the corresponding amplitude or phase reconstruction.

**Fig. 3.**
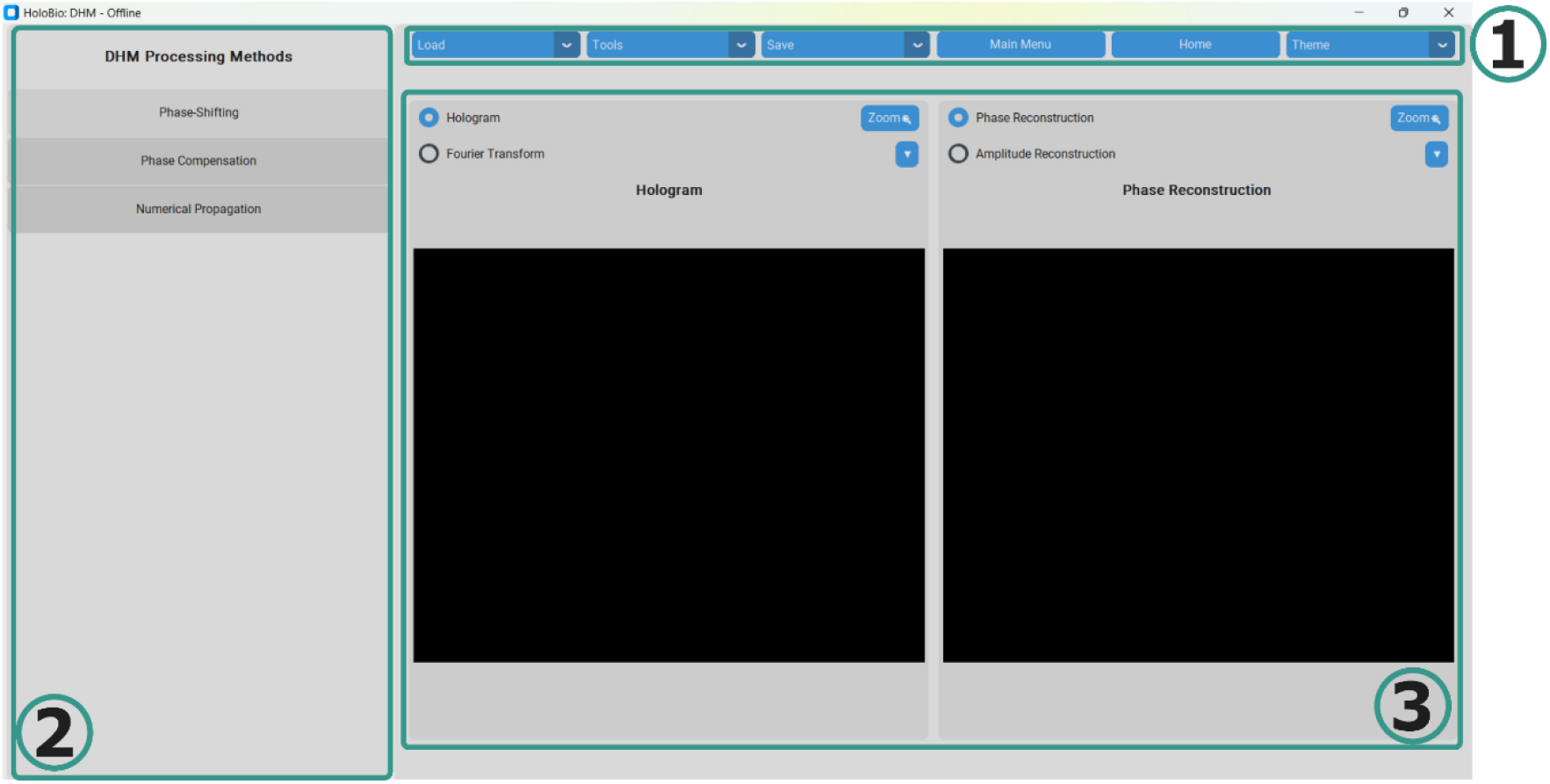
*Offline* DHM package interface in *HoloBio*. The interface is organized into three components: (1) the control panel with core functionalities, (2) the method selection panel, and (3) the visualization panel for hologram and reconstruction display.

#### *Offline* DLHM package

This package provides the same three reconstruction algorithms available in *Real-time* mode: Angular Spectrum, KHDI, and MAS. The interface enables interactive adjustment of the source-to-camera (*L*) and source-to-sample (*z*) distances, thereby facilitating identification of the correct focal plane when the reconstruction distance is unknown a priori. Additionally, users may load a sample-free reference hologram for subtraction from the sample hologram, which significantly reduces background noise and twin-image artifacts in amplitude reconstructions.

Selecting this package opens the interface shown in **Fig. *4***. The interface is also organized into three components. The top control panel (1) contains the primary navigation and management tools, including options to load and save data, access tool menus, return to the main menu or home screen, and change the interface theme. The parameter panel (2) allows the user to configure DLHM-specific reconstruction settings, such as wavelength, pixel pitch in X and Y, magnification, and *z* and *L* distances. Additional controls are available to select the reconstruction algorithm (Angular Spectrum, KHDI, or MAS), define reconstruction limits, and apply hologram reconstruction. The visualization panel (3) displays the selected outputs, enabling users to view the original hologram or its Fourier transform, alongside either reconstructed amplitude or phase images.

**Fig. 4.**
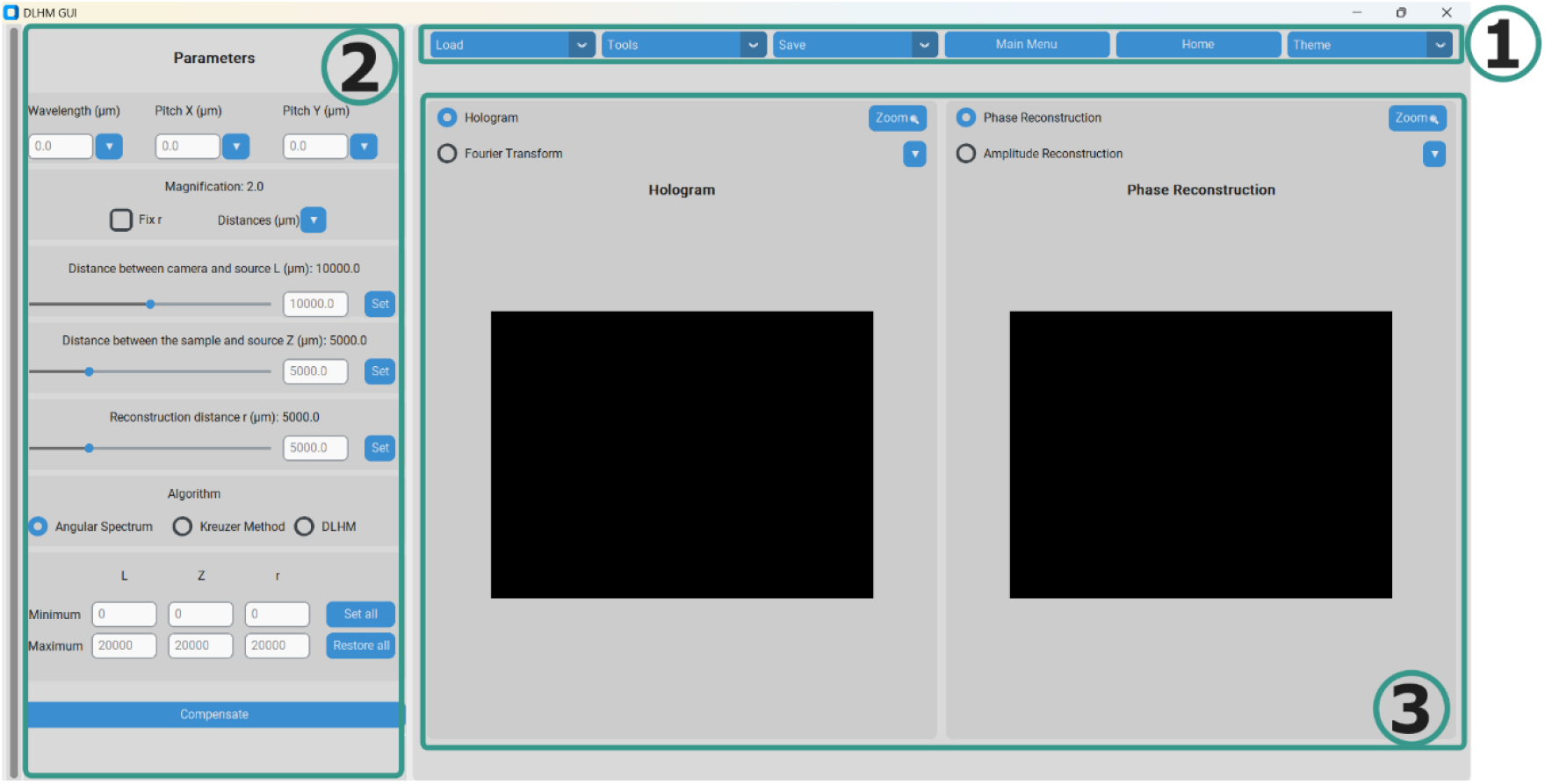
Interface for the *Offline* DLHM package in *HoloBio*. The interface is organized into three components: (1) the control panel with core functionalities, (2) the parameter and algorithm selection panel, where users can set physical parameters (wavelength, pixel pitch, *z* and *L* distances) and choose among available reconstruction methods, and (3) the visualization panel for hologram and reconstruction display.

### Shared Post-Processing and Analysis Tools

For each operational mode, *HoloBio* provides a set of tools that can be grouped into two categories: Processing Toolkits and General-Purpose Analysis functions.

#### Processing Toolkits

- **Bio-Analysis Toolkit**: Designed for biological/biomedical quantification, it allows users to measure phase shifts, extract phase profiles, perform length and thickness measurements, count particles/cells, and estimate sample areas, enabling comprehensive morphological and dimensional analysis.
- **Filters Toolkit**: Enhances image contrast through a variety of spatial filters and colormaps, improving the visualization and interpretation of both amplitude and phase reconstructions.
- **Speckle Toolkit**: Provides tools for the quantification and reduction of speckle noise in selected regions of interest, whether from amplitude, phase, or raw hologram images. Available denoising methods include hybrid median filter (32), mean filter (33), median filter (33), Gaussian filter (33), and SPP filter (34). The toolkit also incorporates comparative visualization features (before/after filtering), speckle reduction plots, and profile extraction to evaluate filtering performance.

The overall architecture of HoloBio, including its operational modes and integrated toolkits, is shown in **Fig. *5***.

**Fig. 5.**
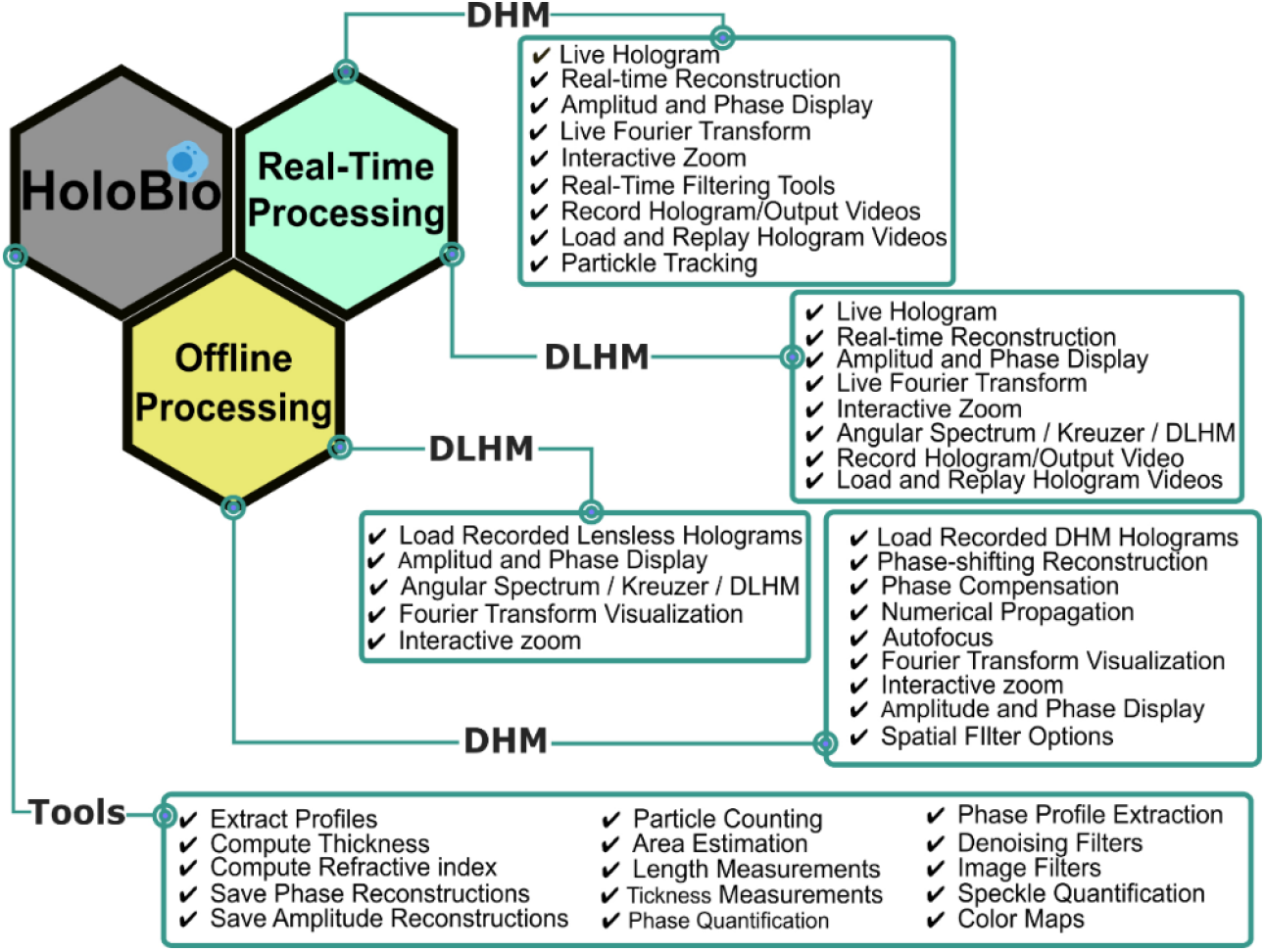
Architecture of HoloBio. The diagram illustrates the two main operational modes: *Real-Time* and *Offline* Processing with their respective DHM/DLHM packages and the integrated toolkits. For a more detailed description of the available tools and configuration options, readers are encouraged to consult the *HoloBio* User Manual.

### Quantitative Biological Imaging with *HoloBio*

This section presents biological imaging applications of *HoloBio*. Each application highlights specific functionalities of the software, demonstrating the utilities of each module and submodule and how the platform supports key tasks such as quantitative phase analysis, cell counting, morphological profiling, and object tracking.

#### Imaging of defocused Red Blood Cell samples

To evaluate *HoloBio* on experimentally defocused off-axis DHM data, we imaged Red Blood Cells (RBC) located at different axial positions from the working distance of the microscope objective (MO) lens. This experiment comprises two steps: (1) numerical reconstruction and phase compensation of the recorded holograms, and (2) autofocusing of the recovered complex wavefield. The RBCs were obtained from Carolina Biological Supply Company (item # C25222) and imaged using a telecentric off-axis DHM system based on a common-path interferometer (35). Illumination was provided by a low-power 532-nm laser diode module (CPS532, Thorlabs). The imaging system includes a 40×/0.75 NA infinity-corrected Nikon MO lens and a 200-mm tube lens. Holograms were recorded using a digital camera with a resolution of 5472 × 3648 pixels and a 2.4-µm pixel pitch. Controlled defocus was introduced by axially translating the sample with a micrometer translation stage.

Using the *Offline* DHM package, the phase compensation process begins by selecting the *Phase Compensation* option, which opens the sub-panel shown in **Fig. *6*A**. This panel is organized into five functional blocks, numbered with Roman numerals for clarity. In section (ii), the user selects the compensation method from the four available options (see Design and Implementation section). These methods are tailored to different optical configurations. For this demonstration, the *Vortex Legendre* fitting method was chosen. Section (iii) allows the user to input the reconstruction parameters, specifically the illumination wavelength and the pixel pitch in both X and Y directions. For this experiment, the values were set to 0.532 µm, 2.4 µm, and 2.4 µm, respectively, according to the experimental setup specifications. Section (iv) contains the Compensation Filter Options, where a spatial filtering method is selected to isolate the +1 diffraction order in the Fourier domain. The *Automatic Circular* filter was selected in this case. Once configured, pressing the *Compensate* button initiates compensation. Upon clicking *Compensate*, a pop-up window displays the filtered Fourier transform of the hologram, as shown in (B), enabling visual confirmation of the selected filter and its position. After closing the window, the software automatically applies the compensation and retrieves the complex object wavefield. The resulting images are shown in **Fig. *6*C-E**. Panel (C) displays the original out-of-focus hologram. (D) shows the initial phase reconstruction (still out of focus), and (E) presents the final numerically refocused phase image. For a detailed description of each compensation method and compensation filter options, readers are referred to the ***HoloBio* user manual**.

**Fig. 6.**
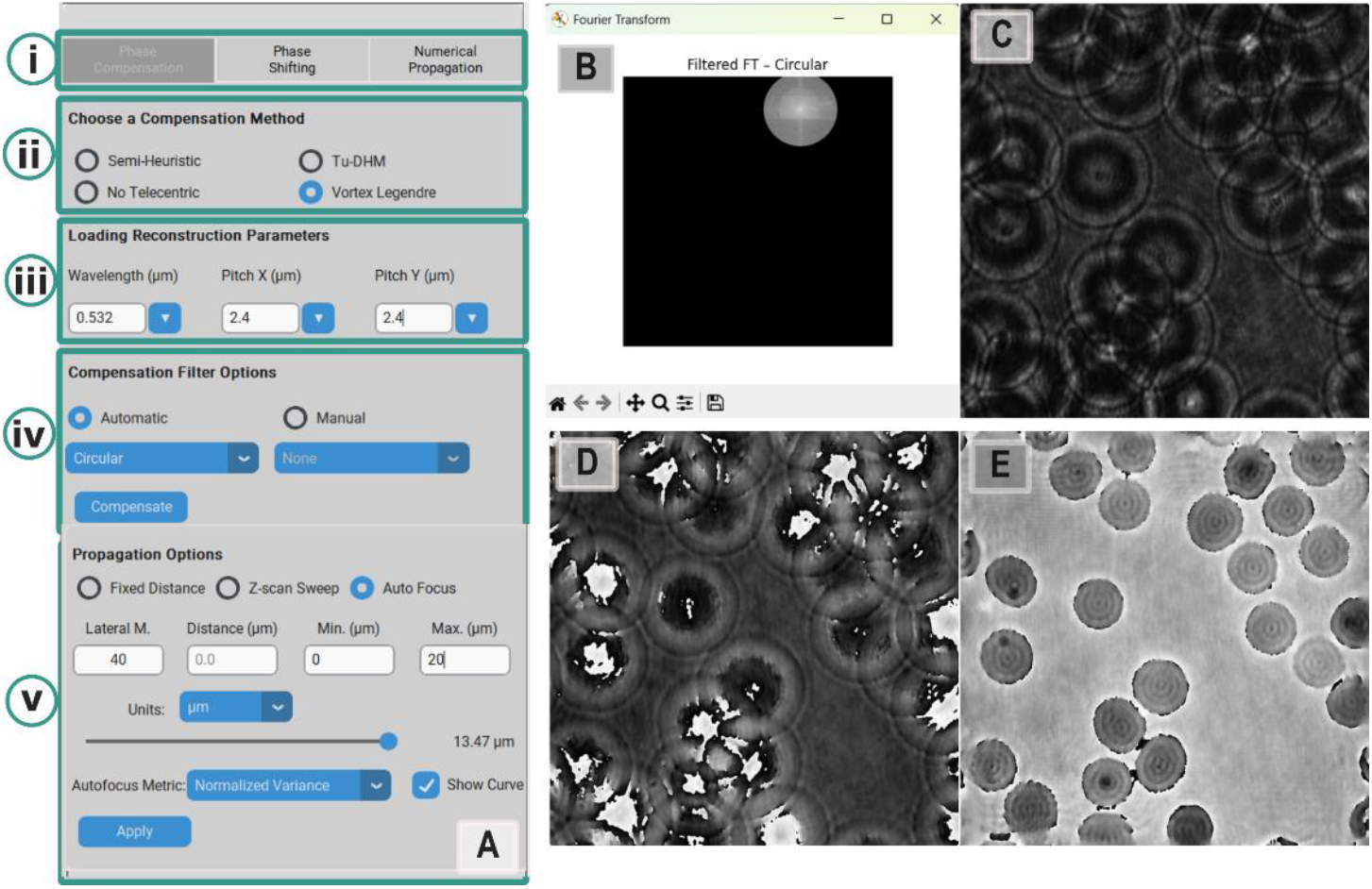
Workflow for phase compensation and numerical refocusing in HoloBio using the Offline DHM Processing package. (A) Interface panel structured into five functional blocks: (i) phase compensation method, (ii) reconstruction parameters, (iii) compensation filter options, and (iv–v) propagation settings. (B) Pop-up window showing the filtered Fourier transform with the isolated +1 diffraction order. (C) Original out-of-focus hologram. (D) Initial phase reconstruction (still out of focus). Finally, (E) numerically refocused phase image after compensation.

In section (iv), users configure the **Propagation Options** subpanel to obtain in-focus images, either in amplitude or in phase. The panel offers three modes: Fixed Distance, Z-scan Sweep, and Auto Focus. In this example, the Auto Focus mode is selected. In this mode, the user defines a minimum and maximum axial propagation range over which the software performs numerical propagation to determine the optimal focus. The optimal focus distance is assessed using one of the two built-in sharpness metrics: Normalized Variance or Tenengrad metric (36). The default metric is the Normalized Variance. Additionally, the user must input the **lateral magnification** of the optical system, in this case 40×, to scale the focus distance correctly to the object space. Once the user clicks the Apply button, a sequence of guided steps is triggered:

- First, a window appears prompting the user to select a Region of Interest (ROI) from the reconstructed image (**Fig. *7*A**). The sharpness metric will be computed exclusively within this selected region.
- After the ROI is selected, a second pop-up window appears, displaying the **Auto-Focus in Progress** indicator (**Fig. *7*B**), while the algorithm evaluates different propagation distances within the specified range.
- If the **“Show Curve”** option is enabled, the software displays the focus curve, which plots the sharpness metric as a function of propagation distance (**Fig. *7*C**). In this example, the minimum of the curve occurs at 13.47 µm, which is identified as the optimal propagation distance (37).

**Fig. 7.**
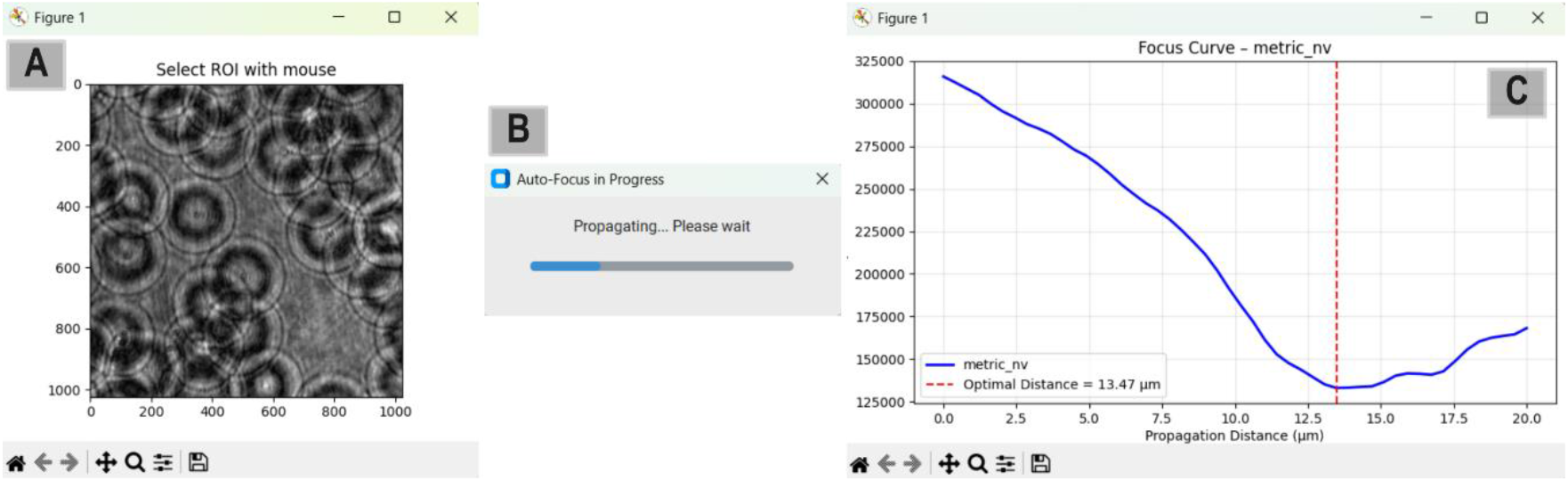
Autofocus workflow in *HoloBio*. (A) A window to select a region of interest (ROI) from the reconstructed image. (B) Auto-Focus progress indicator during propagation. (C) Focus curve showing the propagation distance versus focus metric, with the optimal focal plane indicated.

For additional details on other modes of propagation, propagation algorithms, and autofocus metrics, users are referred to the ***HoloBio* User Manual**.

Once the hologram is properly compensated and focused, the *Bio-Analysis toolkit* provides a set of tools designed for the quantitative assessment of reconstructed amplitude and phase images. As shown in ¡Error! No se encuentra el origen de la referencia. ***Fig*. *8*A**, the panel is divided into three blocks: (i) *Dimensions*, (ii) *QPI Measurements*, and (iii) *Microstructure Metrics*.

#### Dimensions

This block enables users to quantify spatial dimensions within the reconstructed image. First, the user selects the image type to analyze using the radio buttons: *Hologram, Amplitude*, or *Phase*. This choice determines the image on which the dimensional measurement is performed. Next, two key physical parameters must be entered: the *Pixel Size* (in micrometers), which corresponds to the pixel pitch of the digital camera used to acquire the hologram, and the *Lateral Magnification*, which refers to the effective optical magnification applied in the imaging system. In this specific example, a pixel size of 2.4 µm and a lateral magnification of 40× were used. After clicking the *Apply* button, a pop-up window displays the selected image with an overlaid scale bar, as shown in **Fig. 8**. To perform a measurement, the user clicks on two points of interest in the image; a straight line is drawn between the selected positions, and the corresponding distance in micrometers is displayed on screen. In the current example, two RBCs were measured, with zoomed-in regions shown at the bottom for clarity. The results indicate diameters of approximately 8.35 µm and 6.98 µm, respectively. Additionally, the field of view is expressed in micrometers (µm), providing a more intuitive spatial context for interpreting the measurements.

**Fig. 8.**
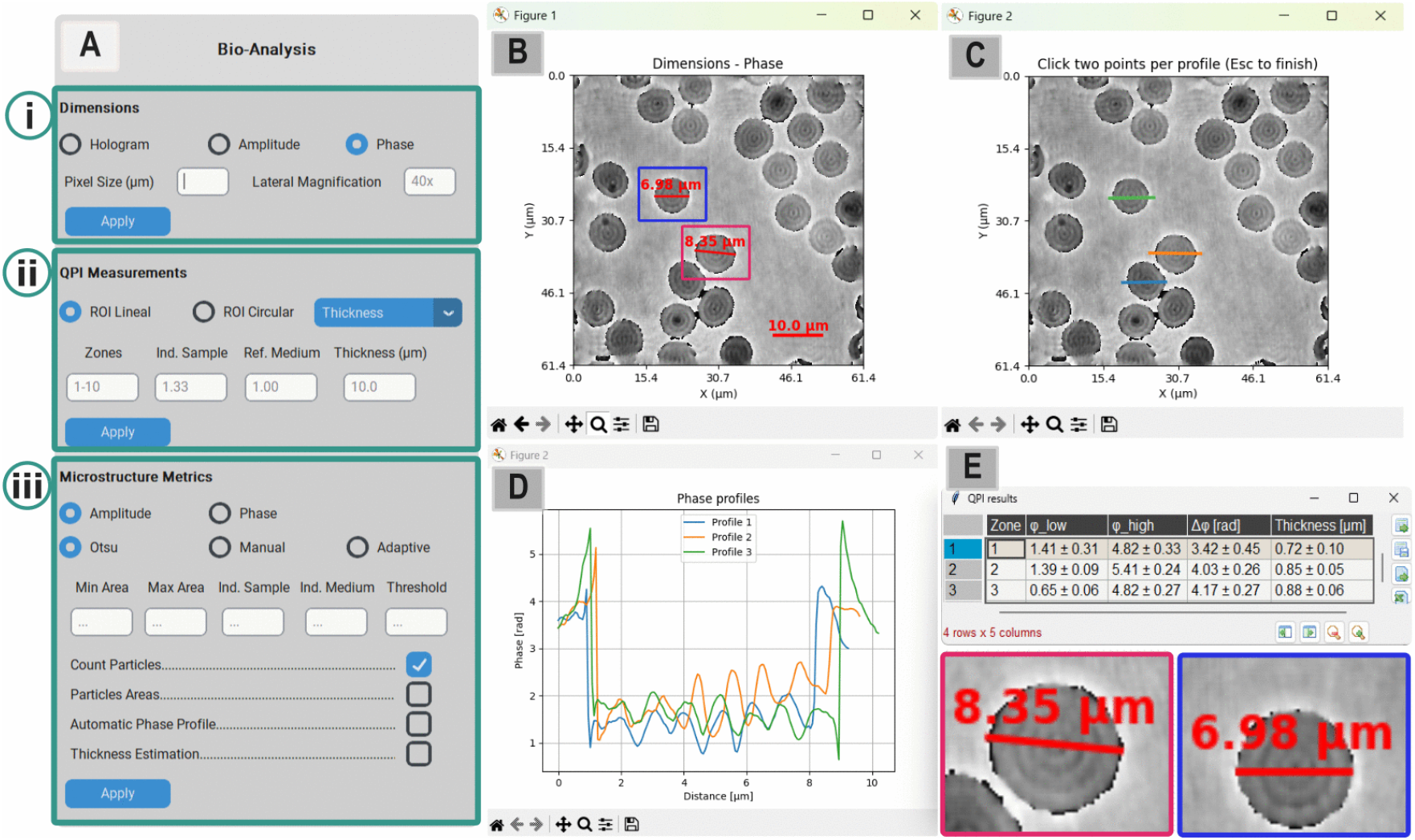
Bio-Analysis workflow in *HoloBio*, showing an example application for dimensional analysis and QPI measurements. (A) Main Bio-Analysis panel with options for dimensions, QPI measurements, and microstructure metrics. (B) Phase image of the sample with dimensional measurements indicated. (C) Window for selecting regions of interest (ROI) directly on the phase image. (D) Phase profiles extracted from the selected ROIs. (E) Summary table of QPI results with corresponding thickness values and example dimensional annotations of the sample.

#### QPI Measurements

This block is designed to perform Quantitative Phase Imaging (QPI) measurements on previously reconstructed phase images. The user begins by selecting the ROI shape using the radio buttons, *ROI Lineal* for straight-line profiles or *ROI Circular* for radial profiles. The dropdown menu on the right allows choosing the measurement mode, either *Thickness* to estimate sample thickness assuming known refractive indices or *Index* to estimate the refractive index assuming known thickness. Once the desired mode is selected, the user enters the physical parameters relevant to the measurement. In this example, we focus on computing the thickness of RBCs, where *Zones* defines the number of phase profiles to be analyzed within the image, *Ind. Sample* refers to the refractive index of the sample, and *Ind. Medium* corresponds to the refractive index of the surrounding medium. For the current analysis, the parameters were set as follows: Zones = 3, Ind. Sample = 1.4 (38), and Ind. Medium = 1.0 (i.e., air). After clicking the *Apply* button, a pop-up window appears allowing the user to define the zones by clicking two points per profile within the phase image (**Fig. 8C**). Once the selection is complete, the software computes the phase profile for each ROI and estimates the thickness based on the provided physical parameters. The results are presented in two output windows, one corresponding to a graphical window showing the phase profiles (**Fig. 8D**), where each profile is plotted in a different color for easy identification; and a results table (**Fig. 8E**) displaying the Zone number, minimum (φ_low) and maximum (φ_high) phase values, phase difference (Δφ), and estimated thickness in micrometers. A summary row at the bottom includes the mean and standard deviation of each column. In this example, the estimated thicknesses for the three selected RBCs are: 0.72 ± 0.31 µm (Profile 1, blue), 0.85 ± 0.05 µm (Profile 2, orange), and 0.88 ± 0.06 µm (Profile 3, green). These results are consistent with typical RBC central thickness values (39), demonstrating the effectiveness of *HoloBio* in performing accurate thickness quantification in biological specimens.

#### Microstructure Metrics

The Microstructure Metrics block provides tools for morphological analysis of biological samples, including *Count Particles, Particle Areas*, and *Thickness Estimation* (***Fig*. *8***-iii). In this case study, the analysis focused on the first two functionalities. Regardless of the selected option, users must choose one of three segmentation strategies: *Otsu, Manual*, or *Adaptive* to isolate relevant microstructures. Additionally, the system allows specification of minimum and maximum area thresholds to filter irrelevant features. For the RBCs sample, this module is used to estimate the projected area of individual cells by selecting the *Phase* image, applying the *Otsu* thresholding method for segmentation, and enabling the *Particles Areas* option. Upon clicking the *Apply* button, three pop-up windows are sequentially displayed: (i) a confirmation dialog asks whether the sample appears white or black in the binary mask, to ensure correct segmentation polarity (**Fig. 9A**); (ii) the resulting binary mask after applying the thresholding method (**Fig. 9B**); and (iii) a histogram of pixel intensities is presented, with the computed threshold overlaid, helping the user to interpret the segmentation result (**Fig. 9D**). After closing the confirmation dialog, a set of three new windows appears. The resulting binary mask (left panel in **Fig. 9C**) shows all connected components detected (19 in total), which are then filtered based on area constraints (Min Area = 8,000 pixels^2^, Max Area = 20,000 pixels^2^). The valid components are outlined and labeled (right panel in C), corresponding to detected RBCs. The Area Analysis Summary (**Fig. 9E**) provides numerical information for each individual particle, including the total number of particles (19), the average area (97.23 ± 13.47 µm^2^), and the area range (74.01–127.68 µm^2^). Additionally, the particle area distribution is shown as a histogram in **Fig. 9F**, illustrating the size variability of the detected RBCs. These measurements fall within the expected range for isolated RBCs [ref], supporting the tool’s accuracy and utility for quantitative analysis. For further information on thresholding options, users are referred to the **HoloBio User Manual**.

**Fig. 9.**
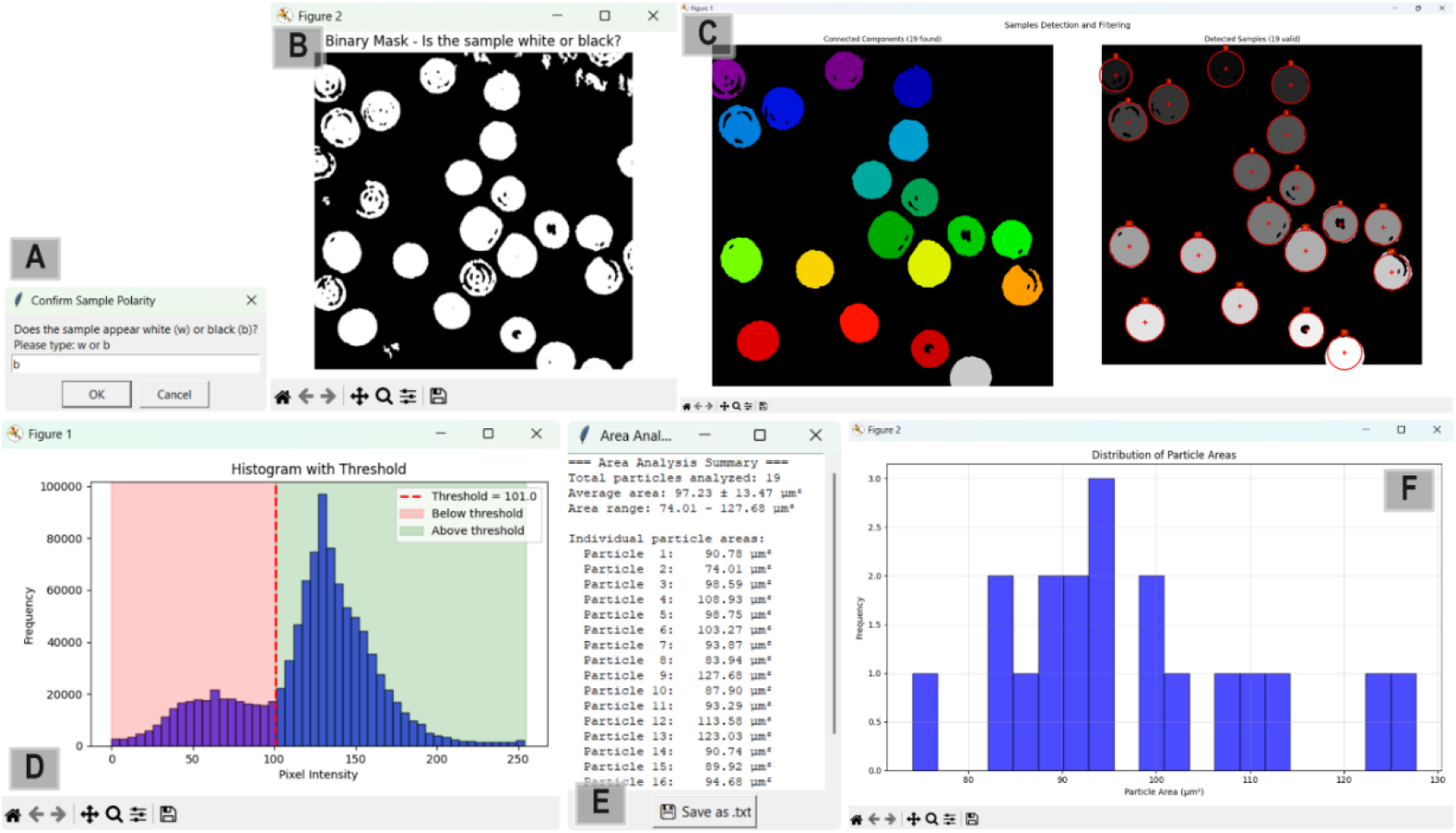
Bio-Analysis workflow in *HoloBio*, showing an example application for particle area estimation. (A) Confirmation dialog for sample polarity. (B) Binary mask obtained after segmentation. (C) Connected components detected (left) and valid particles after filtering by area constraints (right). (D) Histogram of pixel intensities with the computed threshold. (E) Area Analysis Summary window with numerical results of individual particles. (F) Histogram of particle area distribution.

### Speckle noise reduction of onion epidermis tissue samples

An additional biological use case in which *HoloBio* demonstrates its utility for enhancing reconstructed images is speckle noise reduction. As an illustrative example, a set of three phase-shifted holograms of onion epidermis tissue was acquired in a slightly off-axis configuration (40). The reconstruction was performed using the *Offline* DHM package with the *Phase-Shifting* option, specifically the *Blind 3 Raw Frames* method. The reconstruction parameters were set to a wavelength of λ = 632.8 nm and a pixel pitch X and Y of 5.2 µm, providing both amplitude and phase reconstructions suitable for quantitative analysis.

**Fig. 10A** presents the Phase-Shifting interface, highlighting (i) the Offline DHM menu options, (ii) the panel for selecting the phase-shifting method from the available modes (see Design and Implementation section), (iii) the parameter input panel for reconstruction setting, including wavelength and pixel pitch, and (iv) the Propagation Options panel. **Fig. 10B** and **Fig. 10C** show one of the recorded phase-shifted holograms and its corresponding Fourier Spectrum, respectively. The blue rectangles in panel (B) display the two additional phase-shifted holograms used in the reconstruction process. The Fourier Spectrum in panel (C) confirms the slightly off-axis configuration since the spectral orders overlap. The reconstructed amplitude and phase images are shown in panels (D) and (E). respectively, after applying the Blind 3 Raw Frames method.

**Fig. 10.**
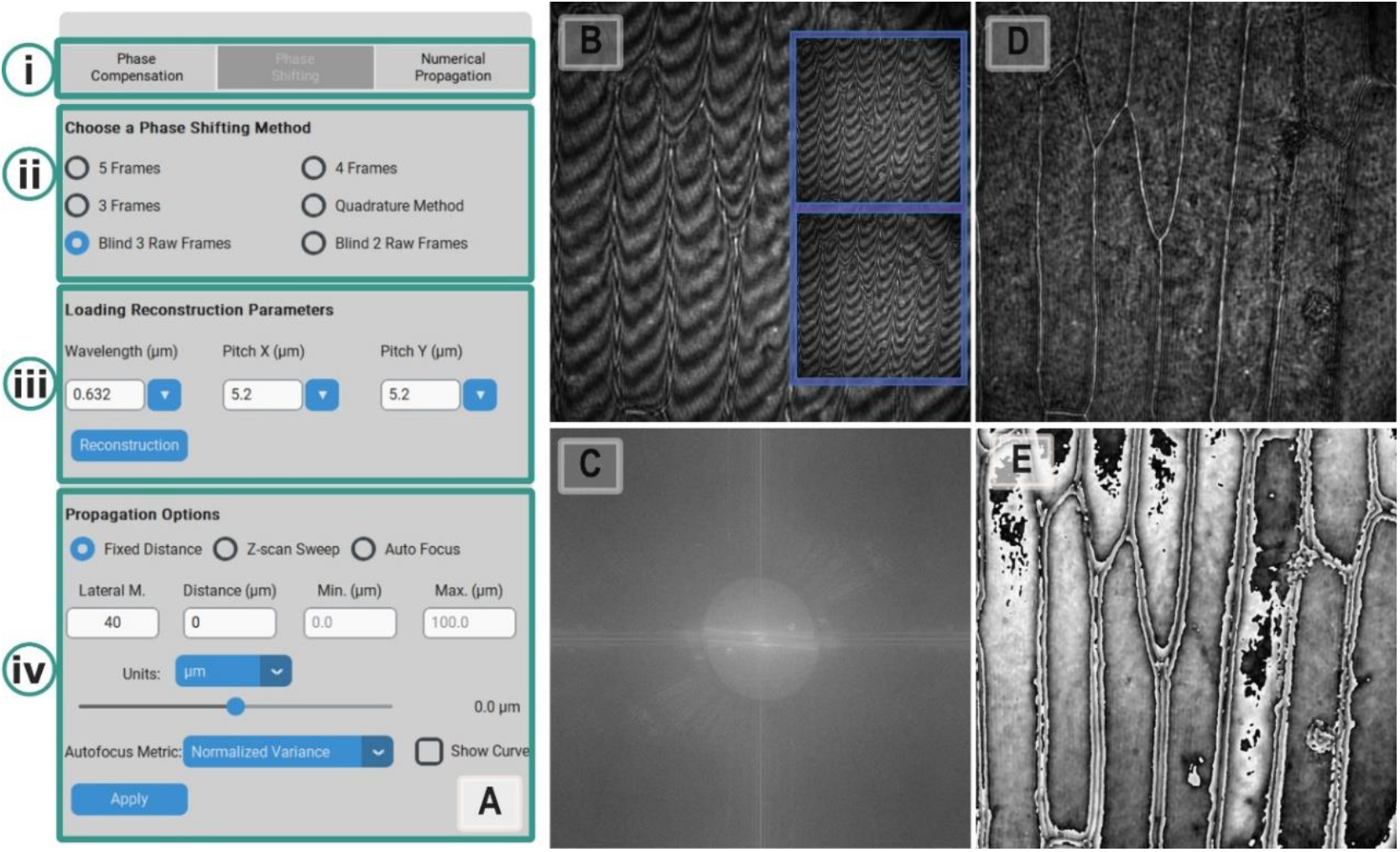
Phase-shifting reconstruction workflow of onion epidermis tissue samples using *HoloBio*. (A) Phase-Shifting interface showing the main menu options, phase-shifting method selection, reconstruction parameters, and propagation options. (B) Example of one recorded hologram, with blue rectangles illustrating fringe shifts. (C) Fourier transform of the hologram, confirming the slightly off-axis configuration. (D) Amplitude reconstruction obtained using the Blind 3 Raw Frames method. (E) Corresponding phase image reconstruction.

Speckle analysis is performed using the *Speckle Toolkit* integrated into *HoloBio*. **Fig. 11A** shows the interface, organized into three functional sections: i) *Speckle Measurements*, where the user selects the image type for analysis (Hologram, Amplitude, or Phase), and defines the number of Zones to be analyzed. Each Zone corresponds to a user-defined rectangular region that is manually selected on the image. For each Zone, the user can further specify the number of Rows and Columns to subdivide the Zone into a mesh of smaller sub-regions. Speckle noise is quantified using the speckle contrast metric, defined as the ratio between the standard deviation and the mean intensity within each sub-region (41). This enables both localized characterization of speckle behavior and the calculation of average speckle metrics over the selected Zones. After setting these parameters, clicking *Apply* opens an interactive window to choose the ROI. ii) *Speckle Filters* provide several speckle reduction methods with configurable parameters. Once the parameters are set, clicking *Apply* applies the selected speckle-reduction method. iii) *Speckle Comparison* offers visualization modes such as *Side-by-Side, Speckle Plot*, or *Profile*. **Fig. 11B** illustrates the Side-by-Side view, enabling direct comparison between the original and denoised images. Two yellow zoom-in regions are shown in both images to highlight local differences in speckle behavior and facilitate visual assessment of the filtering effect. In this example, the amplitude reconstruction of the onion tissue sample is shown after applying the HMM filter with three iterations. **Fig. 11C** displays the selected zones and the generated mesh used to quantify speckle noise. **Fig. 11D** allows quantitative assessment by comparing an intensity profile before and after applying the denoising strategy, showing that the denoised profile is more uniform because of the reduction of the speckle noise. For a detailed description of the Speckle Toolkit and its functionalities, readers are referred to the **HoloBio User Manual**.

**Fig. 11.**
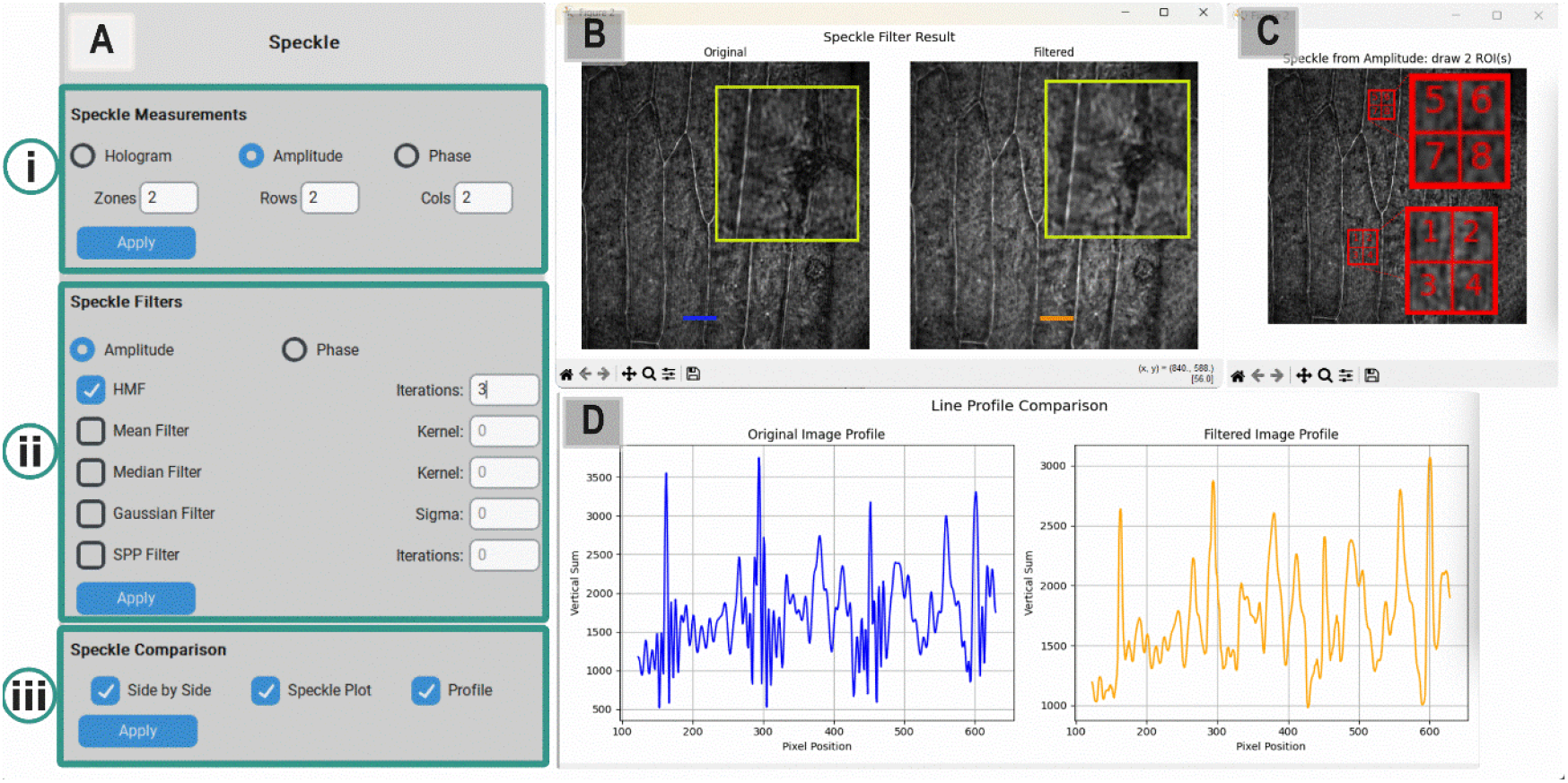
Speckle analysis and reduction workflow in *HoloBio*, illustrating an application of the Speckle Toolkit for a biological sample of onion cell tissue. (A) Interface is organized into three functional sections: Speckle Measurements, Speckle Filters, and Speckle Comparison. (B) Side-by-Side visualization of the original and filtered amplitude reconstructions (using the HMM filter, 3 iterations). (C) Selection of ROIs and subdivision into zones for speckle quantification. (D) Line profile comparison between original and filtered images.

### Wide-Depth Volumetric Imaging of Swimming Paramecia

One of the key advantages of holographic imaging is its ability to recover volumetric information in a plane-by-plane manner from a single recorded two-dimensional hologram. In this application, *HoloBio* was used to process a DLHM hologram (42) of three paramecia swimming in water at different axial positions within an inspection volume of approximately 2–3 mm^3^. **Fig. *12*A** shows the *Offline* DLHM package interface for configuring reconstruction parameters and selecting the reconstruction algorithm. This panel allows manual adjustment of the propagation distance z, enabling each microorganism to be individually brought into focus. **Fig. *12*** shows the reconstructed amplitude images (panels C-E) of the specimens from a recorded hologram (panel A) at three propagation distances. Reconstructed amplitude images were obtained using the KHDI method with the following parameters: wavelength λ = 532 nm, square pixels with a side length of 6.9 μm, distances *L* = 20 mm, and *z* = 6.8 mm (middle paramecium, panel C), 5.6 mm (bottom paramecium, panel D), and 2.8 mm (top paramecium, panel E). Red rectangles indicate zoomed regions of individual paramecia, obtained using the interactive zoom functionality integrated into ***HoloBio***. To further improve visualization of the sample morphology, the **Filters Toolkit** was applied by selecting *Amplitude Viridis* option from the *Filters* menu in the *Tools* bar. A detailed description of the DLHM Offline package and its functionalities is provided in the ***HoloBio* User Manual**.

**Fig. 12.**
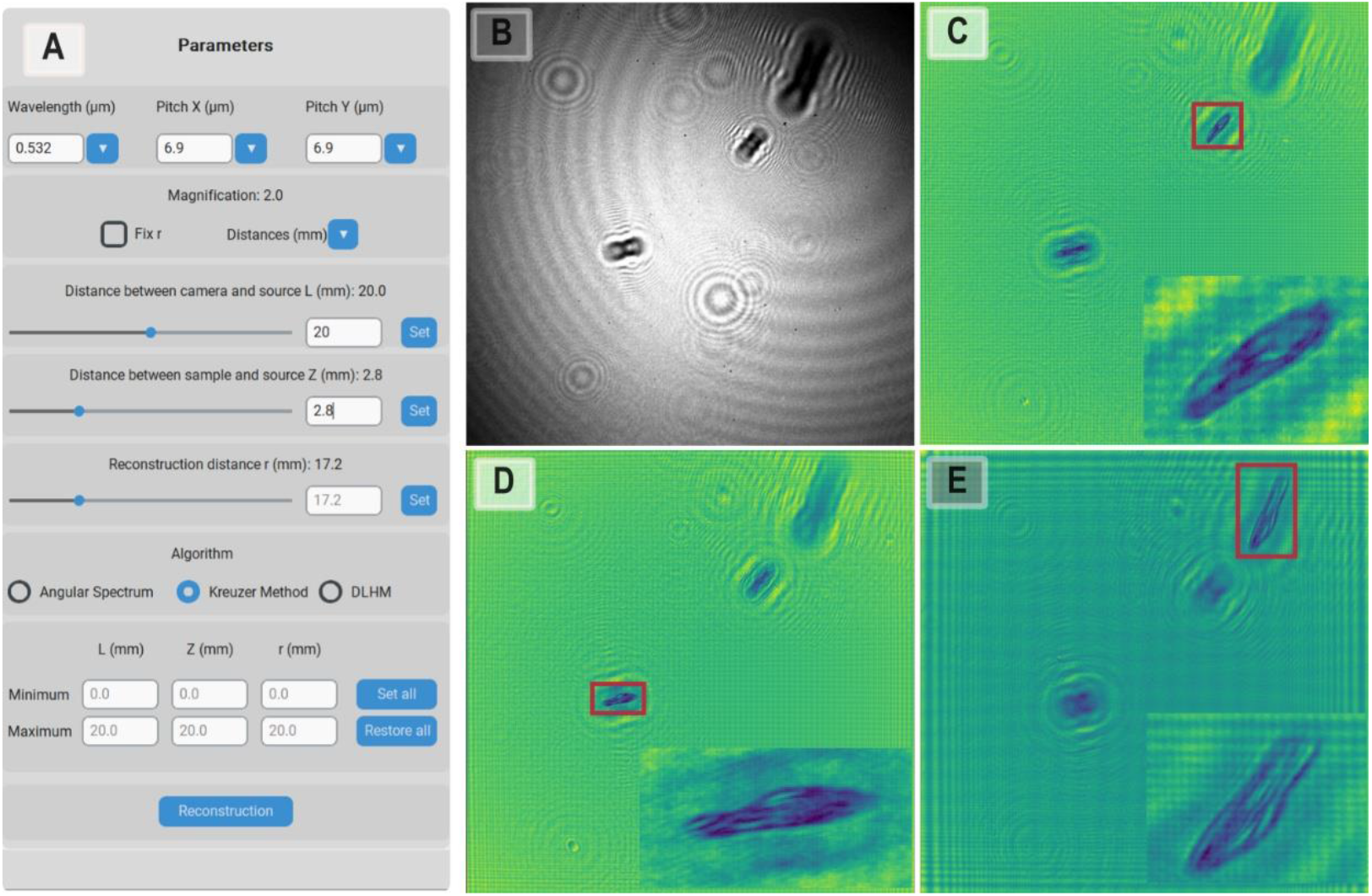
*Offline* DLHM reconstruction of paramecia in water using *HoloBio*. (A) Parameter panel of the *Offline* DLHM package, showing the configuration of reconstruction settings: wavelength λ=532 nm, pixel size 6.9 μm, source-to-camera distance *L*=20 mm. (B) Recorded in-line hologram of paramecia. (C–E) Reconstructed amplitude images at different propagation distances corresponding to individual paramecia. Red rectangles indicate *regions zoomed using HoloBio’s interactive zoom option*.

### Wide-Field Cell Tracking

This experiment was carried out using a previously recorded phase-reconstruction video of RBCs with a duration of *5* seconds at 15 fps. In *HoloBio*, particle tracking begins by uploading the video through **Load > Load Video** from the top control panel. Once loaded, playback can be managed using the **Play** and **Pause** buttons in the *Compensation Control* panel, which allows users to pause at the desired frame to initiate tracking. Before starting, several filters and parameters must be configured in the *Particle Tracking* panel, since the procedure relies on the Kalman algorithm (43); detailed explanations a are provided in the *Tracking Parameters* section in **HoloBio User Manual**. The user may also activate the *World Coordinates* option to report tracking in real spatial units rather than pixel positions. In this example, the phase video was acquired using an off-axis DHM system comprising a 10× microscope objective, and 3.75-µm pixel size digital camera. These parameters where input in the *Parameter* panel. Once all configurations are complete, clicking on the **Tracking** button initiates the analysis, which proceeds as illustrated in **Fig. *13*A–C**. These panels show successive frames where individual cells are detected and tracked. **Fig. *13*D** presents the estimated trajectories in the spatial domain with each path assigned to a unique color and index. **Fig. *13*E** displays the Positions Vector Table containing the frame number, elapsed time, and X–Y coordinates of each tracked sample, thus providing both qualitative visualization and quantitative data for further analysis.

**Fig. 13.**
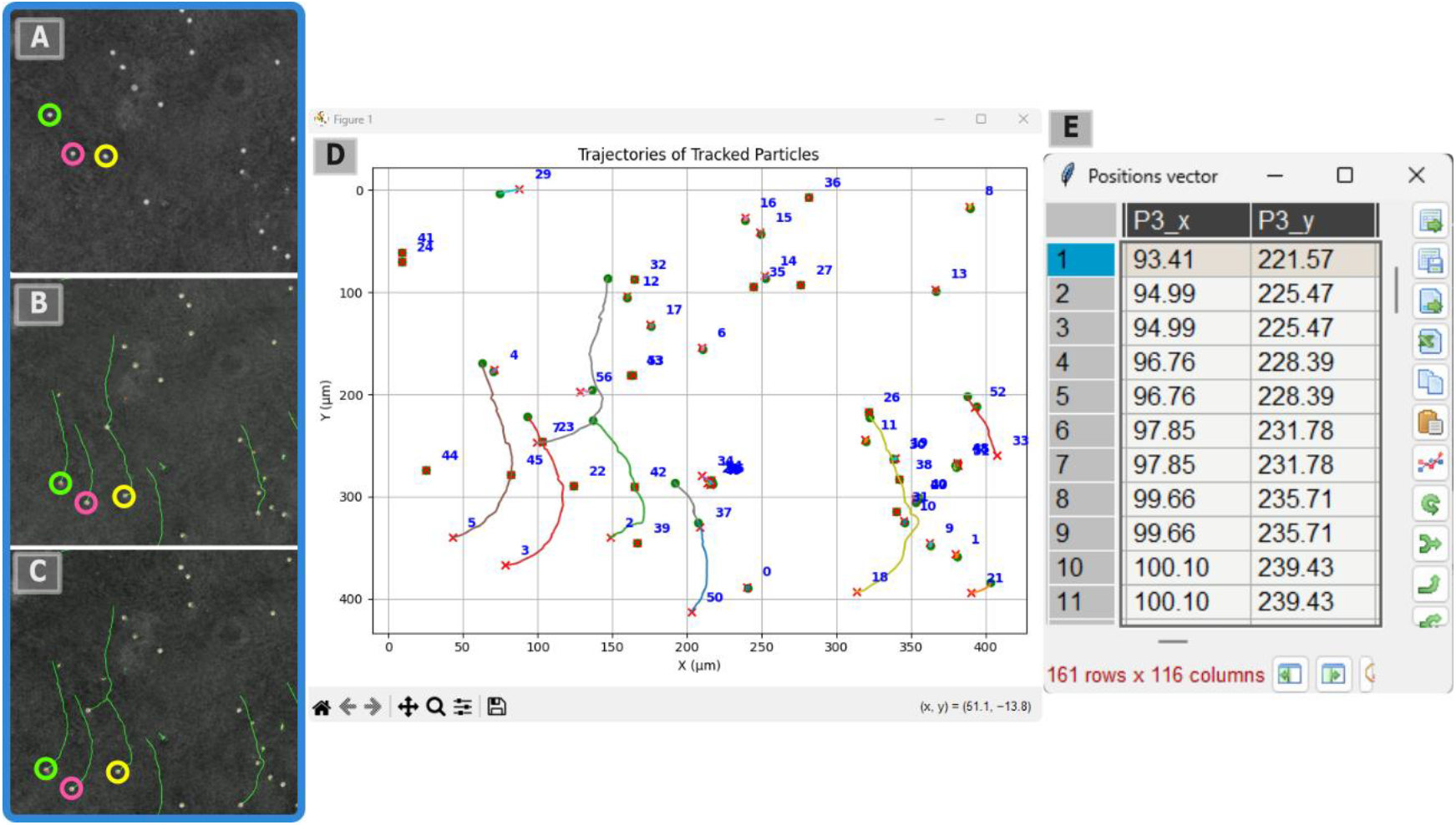
Cell tracking using the *Offline* DHM package in *HoloBio*. (A-C) Consecutive frames from the phase-reconstruction video showing object detection and tracking in real time. (D) Reconstructed trajectories of individual particles in the spatial domain, each displayed with a unique index and color. (E) Position Vector Table summarizing the tracking results, including frame number, elapsed time, and X–Y coordinates of each detected particle.

## Conclusions

This work presents ***HoloBio***, a free and open-source graphical user interface designed to facilitate the use of Digital Holographic Microscopy (DHM) and Digital Lensless Holographic Microscopy (DLHM) in biological imaging. Unlike existing libraries, plugins, or commercial platforms, *HoloBio* provides an integrated environment that unifies hologram acquisition, numerical reconstruction, and biologically oriented analysis tools within a single, intuitive software package. We demonstrated the versatility of *HoloBio* in addressing key tasks relevant to biological research. These included the reconstruction and numerical refocusing of red blood cell holograms, quantitative phase imaging and thickness estimation, cell detection and area measurements, speckle noise quantification and reduction in a complex onion cell tissue, and tracking of cells. A central contribution of *HoloBio* lies in its emphasis on accessibility and usability. By providing modular architecture optimized for real-time and offline processing, and by offering specialized DHM and DLHM packages, the software lowers the technical barriers associated with conventional coding-based libraries and fragmented plugins. Moreover, it has dedicated toolkits, including Bio-Analysis and Filters modules, equipping researchers with traditional functionalities tailored to biological and clinical applications.

*HoloBio* has the potential not only to accelerate the adoption of DHM and DLHM in the life sciences but also to foster reproducibility and innovation in quantitative phase imaging. In this work, HoloBio establishes a flexible and extensible foundation upon which future software enhancements can be built, including the integration of advanced machine-learning models for automated feature detection, support for three-dimensional reconstruction, and expansion toward multi-modal imaging scenarios. Additional development efforts will focus on incorporating GPU-accelerated numerical reconstruction to enable video-rate analysis for functionalities currently available only in offline mode. Furthermore, future updates aim to support real-time particle tracking directly from live holograms, extending HoloBio’s capabilities for high-throughput and dynamic biological analysis.

## Availability

HoloBio-GUI is open-source and is freely available on GitHub https://github.com/SOPHIA-Research-Lab/HoloBio. Users can install it on Linux, macOS, and Windows systems via the Python package manager using the command *pip install holobio-gui*, or by cloning the repository from GitHub. The user manual, usage guide, and video tutorials are accessible in the web page https://sophia-research-lab.github.io/HoloBio/.

## Author Contribution

**W. Mona**: software and validation; **M. Gil, E. Mazo-Gomez, D. Cordoba, R. Restrepo**: software to a lesser extent; **M. Lopera, S. Obando**: validation. **C. Trujillo:** methodology, supervision, the original draft writing, review and editing of the manuscript. **A. Doblas:** methodology, supervision, validation, manuscript review and editing, and funding acquisition. **R. Castaneda**: software, methodology, supervision, validation, writing of the original draft, and review and editing of the manuscript.

## Acknowledgments

The authors also express their gratitude to all previous contributors and collaborators whose prior work laid the groundwork for the present work, although they were not directly involved in the development of *HoloBio*. W. Mona, M. Gil, C. Trujillo, and R. Castaneda acknowledge funding from the Vicerrectoría de Ciencia, Tecnología e Innovación, Universidad EAFIT. A. Doblas acknowledges the support provided by National Science Foundation (2042563 and 2404769).

